# Entropy, complexity, and maturity in children’s neural responses during naturalistic mathematics learning

**DOI:** 10.1101/2020.11.18.387431

**Authors:** Marie Amalric, Jessica F. Cantlon

## Abstract

A major goal of human neuroscience is to understand how the brain functions in the real world, and to measure neural processes under naturalistic conditions that are more ecologically valid than traditional laboratory tasks. A critical step toward this goal is understanding how neural activity during real world naturalistic tasks relates to neural activity in more traditional laboratory tasks. In the present study, we used intersubject correlations to locate reliable stimulus-driven neural processes among children and adults in naturalistic and laboratory versions of a mathematics task that shared the same content. We show that relative to a control condition with grammatical content, naturalistic and simplified mathematics tasks evoked overlapping activation within brain regions previously associated with math semantics. We further examined the temporal properties of children’s neural responses during the naturalistic and laboratory tasks to determine whether temporal patterns of neural activity change over development, or dissociate based on semantic or task content. We introduce a rather novel measure, not yet used in fMRI studies of child learning: neural multiscale entropy. In addition to showing new evidence of naturalistic mathematics processing in the developing brain, we show that neural maturity and neural entropy are two independent but complementary markers of functional brain development. We discuss the implications of these results for the development of neural complexity in children.

## Introduction

Children typically learn from educational narratives in which new information builds on past information over the course of seconds to minutes. In contrast, typical laboratory tasks present isolated stimuli repetitively at intervals of milliseconds to seconds. Repetitive, fast-paced, controlled paradigms are important for making pure contrasts to test hypotheses but they may be missing important aspects of context or continuity that are critical for children’s computations (Cantlon, 2020). Moreover, the natural pace of learning is often absent in traditional neuroimaging paradigms. EEG and fMRI studies typically analyze neural signals over intervals of milliseconds to a few seconds. Slow fluctuations in neural signals are often considered artifactual noise and neutralized. That approach neglects neural processes that accumulate knowledge, meaning, or information over time (Hasson et al., 2015). In this sense, overly simplistic neuroimaging paradigms could neutralize, ignore, or remove neural processes that are critical to human cognition and development. An alternative approach is to use naturalistic tasks in which information gradually unfolds in educational narratives and examine the temporal fluctuations in neural signals across timescales.

Recent neuroimaging studies have tested children with naturalistic educational videos to measure the similarities between neural processes in children and adults (Cantlon and Li, 2013; Emerson and Cantlon, 2015; Kersey et al., 2019; Lerner et al., 2019; Richardson et al., 2018). Those studies use intersubject correlation measures to quantify the similarity in neural timecourses between children and adults. In intersubject correlation, the neural timecourse at each voxel is tested for temporal correlation among subjects. Significant correlation in neural responses among subjects indicates the presence of a reliable stimulus-driven neural process in that brain region. These studies established that naturalistic paradigms evoke patterns of neural activation that associate or dissociate by function and age in predictable and reliable ways. For example, Cantlon and Li (2013) showed that 4-year-old children exhibited temporal correlation in the intraparietal cortex when they were watching number-related “Sesame Street” videos, while letter-related videos elicited correlated neural responses in Broca’s area. Furthermore, children’s neural similarity with adults in the parietal cortex predicted their performance in mathematical tests, while their neural similarity with adults in Broca’s area predicted their performance in verbal tests. Kersey et al. (2019) confirmed the functional dissociation between math-versus reading-related natural viewing activity, and demonstrated that both math- and reading-related regions show immature neural patterns in 4 to 8-year-old children. However, in prior naturalistic studies of children it is unclear how neural activity during naturalistic tasks is similar to or different from activity in the more traditional simplified tasks because they were not directly compared. This question is important in the study of child development because, as mentioned earlier, critical features of the pace and context of learning may be unique to naturalistic tasks (Cantlon, 2020).

Here we compare naturalistic and simplified versions of a mathematics task which presents common content. In the naturalistic task a teacher explains formal arithmetic principles in an educational narrative whereas in the simplified task the principles are queried in a traditional two-alternative forced choice task. The core prediction is that the naturalistic and simplified versions of the arithmetic tasks should evoke overlapping activation within brain regions that process the common semantic content of the tasks, relative to a naturalistic control condition testing non-mathematical content (a grammar lesson). After testing this hypothesis, we explore novel techniques to compare the temporal signatures of the naturalistic and simplified tasks and examine neural differences in those processes. One measure that has not been used to study brain development that could be useful in the study of children’s learning is the temporal entropy of the neural signal (Vanderwal et al., 2019). Entropy classically describes the degree of organization of a time series. In other words, it provides a measure of the complexity of the underlying structure of a time series, and evaluates its degree of unpredictability. For example, consider one series with alternating 0 and 1, and another series with values either 0 or 1 randomly chosen with probability 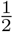. These two series have the same mean and variance, yet their entropy differs greatly. In the first one, each value can be predicted on the sole basis of the previous one, the series is thus highly regular and has low entropy. On the contrary, values of the second series are unpredictable, and that translates into high entropy. Now consider a series composed of a repetitive pattern made of 0s and 1s (for example [0 0 0 1 1 0 1 0 0 0 1 1 0 1 0 0 0 1 1 0 1 ...]). Each value in this example repeats after 7 occurrences, this series thus has a certain degree of structure that makes it somewhat regular. Its entropy falls between the entropy of the two time series above.

The Kolmogorov complexity is an exact measure of this degree of organization (Grunwald and Vitányi, 2004). However, for physiological data that only contains a moderate number of time points, it can only be approximated. Several approximate measures of the Kolmogorov complexity have been developed such as sample entropy (SampEn) (Richman and Moorman, 2000). In the past decades, it has been combined with the method of multiscale entropy (MSE) to analyze the complexity of biological signals (Costa et al., 2005, 2002). This method has first been applied to many types of biological signals such as human gait (M. Costa et al., 2003) or heart rate (Norris et al., 2008), and to electrophysiological data (Heisz and McIntosh, 2013). The evaluation of multi-scale entropy of EEG signals has particularly proven useful to characterize the severity of Alzheimer’s disease (Azami et al., 2017; Escudero et al., 2006; Nobukawa et al., 2019; Park et al., 2007; Yang et al., 2013b), or autism spectrum disorder risk (Bosl et al., 2011; Catarino et al., 2011), but also as a marker of stress (Ahammed and Ahmed, 2020) or as an indicator of the automatization of cognitive processes (Grundy et al., 2017; Wang et al., 2019)

While the literature is rich in application of the evaluation of multiscale entropy of EEG signals, it has only recently extended to the fMRI literature. In this technique, the sample entropy of the BOLD signal is evaluated over various timescales in each voxel of the brain. At scale 1, the sample entropy of the full BOLD signal is evaluated. In a given voxel, if the BOLD signal follows a regular simple pattern, the entropy value of this voxel is low, whereas it is higher if the BOLD signal follows a regular but complex pattern, and even higher if the BOLD signal varies unpredictably over time. At other scales, it is the sample entropy of coarse-grained signals that is evaluated. For example, each data point of the “scale 10 time series” is calculated as the mean of 10 consecutive and non overlapping data points of the initial series, such that the length of the “scale 10 time series” is one tenth of the length of the initial series (see Figure 1). The method of multiscale entropy thus reveals the dynamics properties of the BOLD signal over a wide range of timescales.

Although entropy is thought of as disorder in thermodynamics (Boltzmann, 1964), some evidence indicates that increased entropy in information processing and neural signaling reflects advanced processing. For example, neural entropy is positively correlated with intelligence test performance in adults (Saxe et al., 2018; Yang et al., 2013a). Thus, on the one hand, entropy is considered a marker of disorder and randomness in material sciences. On the other hand, entropy is interpreted not as disorder but as richness in cognitive and neural processing. Here we compare entropy over time in the neural signals generated by children and adults during naturalistic and controlled learning tasks. If neural entropy reflects the integration of information over time then entropy should be greater in the naturalistic learning condition compared to the controlled task over a longer period of time. Moreover, if entropy is a measure of advanced neural processing then entropy should increase with development.

**Figure 1:**
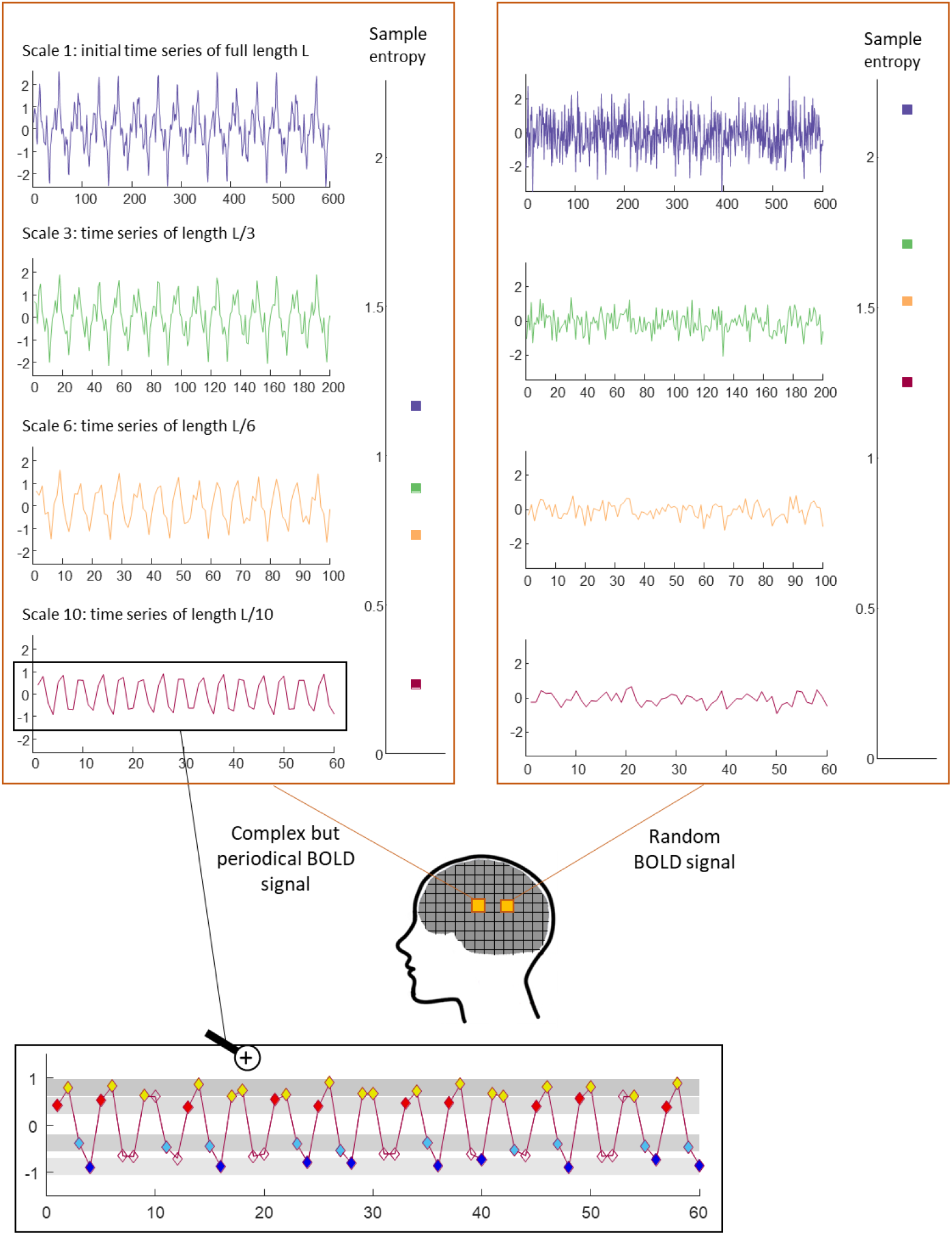
Multiscale entropy. *Top*: In each voxel of the brain, determination of multiscale entropy for a periodical BOLD signal (left) and a random BOLD signal (right). Each data point of the scale n signal is created by averaging n consecutive non-overlapping data points of the original timeseries, thus resulting in a timeseries of length L/n of which the sample entropy is calculated. Entropy values in the case of a random signal are higher than in the case of a periodical signal. *Bottom*: Zoomed in illustration of the method to calculate sample entropy: first evaluate the number of repetitions of each pattern of length two throughout the timeseries, within a given tolerance range. In this example, the pattern “red-yellow” repeats 9 times, the pattern “yellow-blue” repeats 8 times, the pattern “blue-blue” repeats 7 times and the pattern “blue-red” repeats 4 times, for a total of 28. Then evaluate the number of repetitions of each pattern of length three throughout the timeseries, within the same tolerance range. Here, the pattern “red-yellow-blue” repeats 6 times, the pattern “yellow-blue-blue” repeats 7 times, and the pattern “blue-blue-red” repeats 4 times, for a total of 17. Finally calculate the negative log ratio of these two numbers: −log(17/28) = 0.217.

## Results

### Method

32 participants (14 adults and 18 third-grade children) were scanned during fMRI while completing a two-alternative forced choice math task, a naturalistic mathematics education lesson, and a naturalistic grammar education lesson. Both the controlled math task and the naturalistic math video lesson shared the same mathematical content. The forced-choice task directly tested participants’ understanding of the commutative principle of multiplication. On each trial, they were given a few seconds to decide whether two symbolic operations or two dot arrays, including commutative pairs (e.g. “2×3” versus “3×2”, see Table S1 for a full list of stimuli), were numerically equal or not (top row of Figure 2). The math video was a lesson describing and explaining the commutative principle of multiplication. While the controlled math task and math video shared content about the commutative principle, the grammar video and math video, in contrast, shared superficial features of a naturalistic educational lesson. The content of the grammar lesson was the principle of combining relative pronouns into phrases. Both the math and grammar videos were designed to parallel a real world school lesson as much as possible by presenting formal information in a building narrative. Both lessons included the same virtual teacher explaining either a mathematical or grammatical rule over three 80-second video sequences that allowed a natural progression in the lesson from a simple example, to a more complex example, to a general definition and application of the principle.

**Figure 2:**
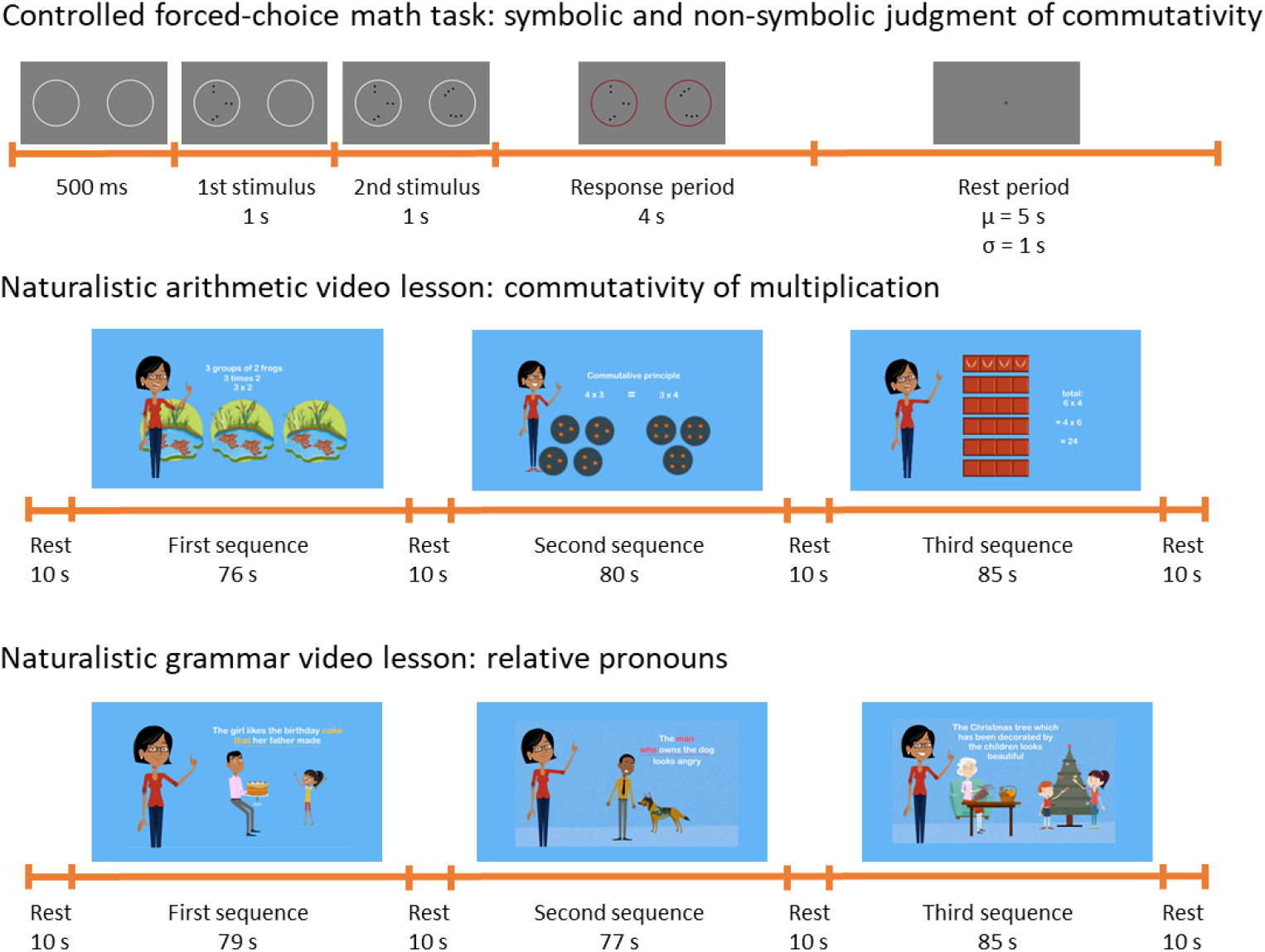
Protocol. *Top*: Timeline of a trial of the two-alternative-forced-choice math task. Participants were given 4 seconds to decide whether the results of two simple operations or the numbers of dots in two different sets were the same or were different. *Middle and bottom*: Timeline of the presentation of both arithmetic and grammar educational videos and representative frames extracted from each of the three sequences composing the videos.

### Behavioral results in the controlled same/different math task

Children correctly answered 89.0 ±1.84% of the trials. Children’s accuracy did not significantly differ from adults’ accuracy (92.4 ± 2.69%, t(30) = 1.11, p = 0.275), although they generally answered slower than adults (children: 1.97 ± 0.077s; adults: 1.08 ± 0.108s; t(30) = 7.03, p < 0.001). Children failed to respond on 13.7% of the trials while adults only missed 1.98% of trials. When counting missed trials as errors, children reached an overall accuracy of 76.9 ± 2.43%, significantly lower than adults’ accuracy (90.8 ± 2.98%, t(30) = 3.58, p < 0.002). Whether considering missed trials or not, children performed significantly better than chance for each condition (symbolic/non symbolic × 3 categories of pairs; all ps < 0.05), and so did adults (all ps < 0.001).

### fMRI results

#### Mathematics and grammar tasks yield dissociable neural responses during naturalistic learning

We first searched for cerebral activity exhibiting higher intersubject correlations (ISC) during the viewing of the arithmetic video lesson than during the viewing of the grammar one. For both children and adults the resulting map revealed extensive correlated activity in parietal areas and in the right inferior frontal gyrus (left panels of Figure 3). Among adults, additional correlated activity was found in the bilateral posterior inferior temporal gyri (middle left panel of Figure 3). Among children, we note that the arithmetic video elicited more correlated activity than the grammar video in regions very similar to adults, mostly in the right intraparietal sulcus and the right inferior frontal gyrus pars opercularis (bottom left panel of Figure 3).

**Figure 3:**
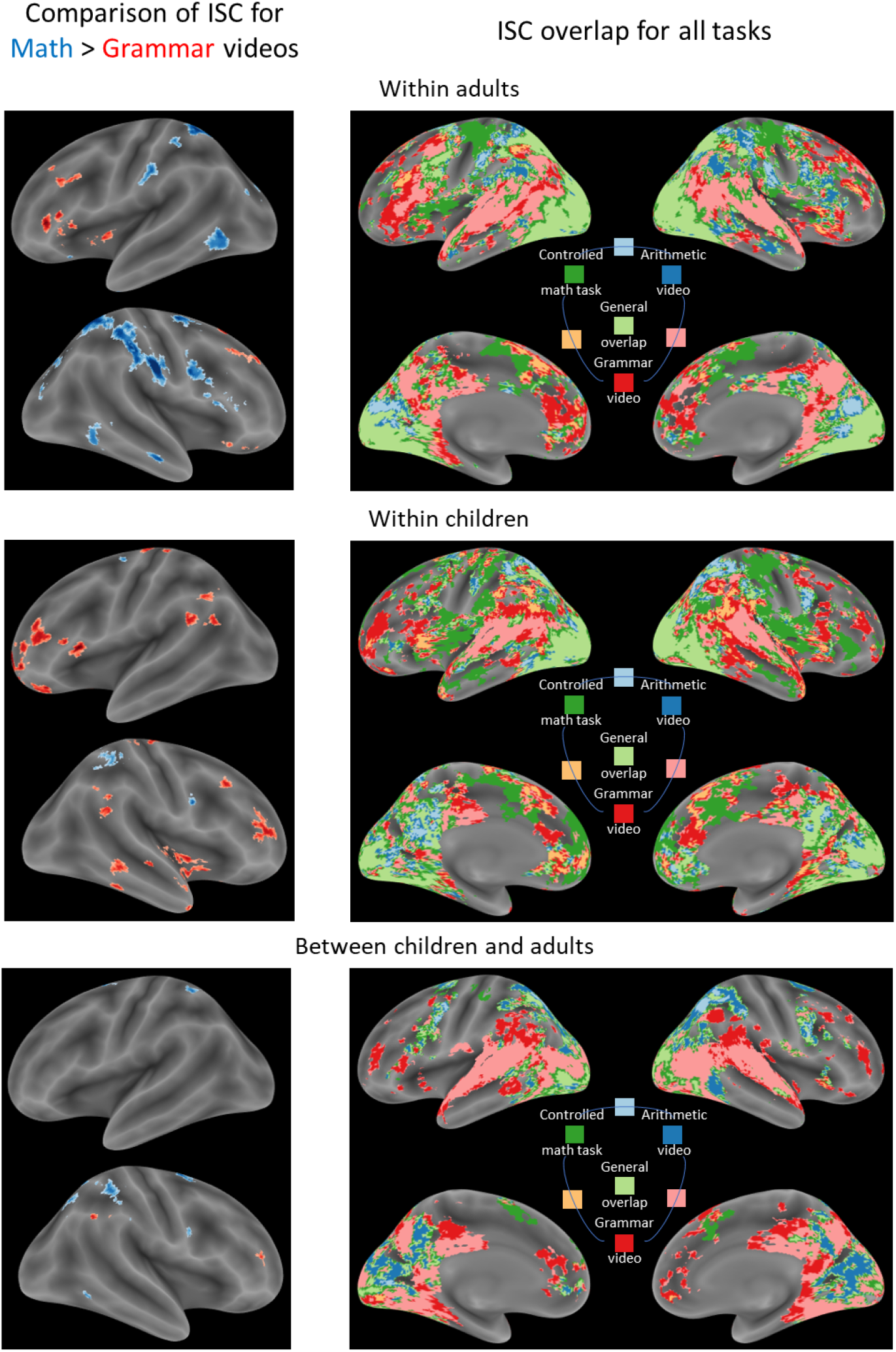
Correlated activation during naturalistic and controlled tasks reveals both modality and content preferences. Left: Comparison of intersubject correlations for arithmetic (in blue) versus grammar (in red) educational videos (top) among adults, (middle) among children, and (bottom) between children and adults. *Right*: Maps showing the significant correlations of neural timecourse during each task, and their respective overlaps. In particular, red, dark blue and dark green clusters reveal correlated activation respectively associated with each one of our three tasks. Light blue clusters represent correlated activation that is common to both math tasks, while pink clusters represent correlated activation that is common to the viewing of both educational videos. Finally, light green clusters locate the brain regions exhibiting significantly correlated activation for all three tasks together. All maps are displayed at p < 0.001 voxel-wise and FDR p < 0.05 cluster-wise.

The converse contrast of grammar versus arithmetic video lessons yielded intersubject correlations in inferior frontal regions for both children and adults (left panels of Figure 3). In adults, such clusters were mostly left-lateralized in the inferior frontal gyrus pars triangularis and pars orbitalis. Children however, exhibited more bilateral grammar-related activity in the middle frontal gyri, and at the temporo-parietal junction (note that this lateralization difference between children and adults was confirmed by a sensitive single-subject region of interest (ROI) analysis detailed in Supplementary Materials).

### Naturalistic mathematics learning and the controlled mathematics task show overlapping neural responses in the parietal cortex

We next compared the intersubject correlations of neural responses elicited by the controlled arithmetic task to those from the naturalistic arithmetic video lesson. We observed a remarkable overlap of correlated activity for both mathematical tasks in the bilateral parietal cortex, and the right inferior frontal gyrus (right panels of Figure 3) in both children and adults, and additionally in the bilateral posterior inferior temporal gyrus in adults. The main activation differences between tasks were in the motor cortex for the controlled math task (participants answered by pressing buttons), and activity along the superior temporal sulcus for the naturalistic arithmetic video lesson, noticeably overlapping neural responses during the grammar video lesson, as both videos included spoken language (see Supplementary Materials for a complete description of regions exhibiting overlap between our three tasks).

To quantitatively assess the spatial overlap of the neural responses elicited by both naturalistic and controlled math tasks, we performed a sensitive single-subject representational similarity analysis in five main math-related ROIs (bilateral IPS, bilateral pITG, and right IFG Oper). For each subject, across all voxels of our ROIs, we computed the correlation coefficients between the ISC values observed during each pedagogical video as well as during the forced-choice math task. We then compared the correlation of ISC values corresponding to the two math tasks versus the correlation of ISC values corresponding to the two videos (paired t-tests). In children and adults, intersubject correlations in the left and right IPS were more spatially correlated between the naturalistic arithmetic video and the controlled arithmetic task than between the naturalistic arithmetic and grammar videos (ts(31) > 4.08, ps < 2.10-4; note that to reach significance after Bonferroni correction for multiple comparisons in 5 ROIs, the p value must be less than 0.01; also note that this result was true for children and adults separately, all ps < 0.006). A smaller but nonetheless significant difference was also found in right pITG (t(31) = 2.78, ps = 0.0047).

### Naturalistic tasks elicit greater entropy than controlled tasks, particularly at longer times-cales

We then turned to the comparison of the dynamic signatures of neural activity during the naturalistic versus controlled tasks. To do so, we computed the entropy values in each voxel of the brain over 10 timescales (method of multiscale entropy, see Figure 1 and details in the Methods section). We then averaged the entropy maps of adults and children over short timescales (1 to 5) on the one hand, and over long timescales (6 to 10) on the other hand, separately for the arithmetic and grammar video lessons, and for the controlled math task. To evaluate which brain regions exhibited entropy differences between the naturalistic and controlled tasks, we entered these resulting mean entropy maps in a general ANOVA. At short timescales, we did not observe any significant difference between the videos and the math test in children. In adults, the naturalistic math task elicited greater entropy than the controlled task in the occipital cortex, the cuneus, and the right fusiform gyrus. At longer timescales, multiple sites in the occipital cortex, the inferior and superior temporal gyri, the parietal cortex and the inferior frontal gyrus distinguished the naturalistic and controlled tasks for children and adults alike. Sites where the arithmetic video lesson elicited greater entropy than the controlled math task are represented on Figure 4 and listed in table S2. We note that these maps were remarkably similar between children and adults, except in the occipital cortex, where adults showed a greater entropy difference compared to children across timescales.

**Figure 4:**
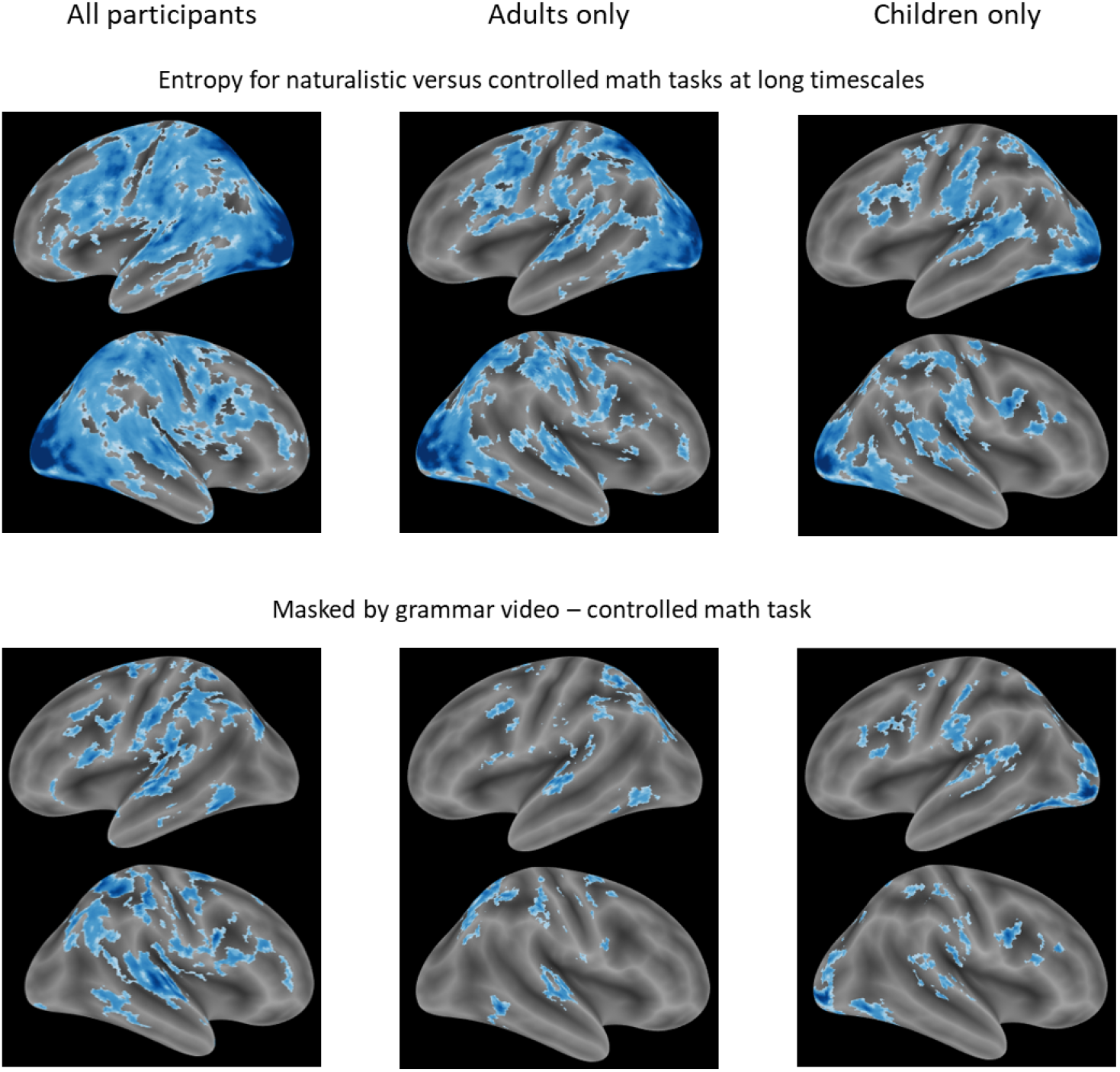
Dynamic differences in modality and content at long timescales. *Top - Difference in modality*: Maps of contrast of long timescales (6 to 10) entropy between the naturalistic and controlled math tasks in all participants together (left), adults (middle), and children (right). *Bottom - Difference in content*: Maps of the remaining clusters after masking by the contrast of long timescales entropy between the grammar video and the controlled math task. All maps are displayed after a 2 mm FWHM Gaussian smoothing, with p < 0.001 voxel-wise and FDR p < 0.05 cluster-wise.

We then tested whether long scale neural entropy was modulated specifically by mathematical content. The whole-brain direct comparison of math versus grammar video lessons with strict correction for multiple voxel comparisons did not reveal any significant differences in entropy. However, a more sensitive apriori ROI analysis showed that the math video lesson elicited greater entropy than the grammar video lesson, particularly in bilateral IPS, left pITG, and right IFG Oper (ts(31) > 1.93, ps < 0.031). We then confirmed this content preference in a whole-brain analysis by masking the contrast of arithmetic video lesson versus the controlled math task by the entropy corresponding to the grammar video lesson in a contrast with the controlled math task (bottom row of Figure 4). In adults, this map included bilateral parietal sites, bilateral inferior temporal sites, and the bilateral inferior frontal gyrus pars opercularis. In children, sites where the modulation of entropy was particularly sensitive to mathematical content were mostly observed bilaterally in the inferior frontal gyrus, as well as in the right superior parietal lobule (see table S3 for full lists). Taken together, these results suggest that over long periods of time, the entropy of the neural signal in the parietal cortex was not only enhanced by the type of task (naturalistic more than controlled), but also by the semantic content of the task (math more than grammar).

### Functional connectivity of parietal regions is enhanced during naturalistic tasks compared to controlled tasks

Higher long scale neural entropy in the parietal cortex for naturalistic versus controlled math tasks could be the result of long-range interactions between parietal regions and other regions of the brain (McIntosh et al., 2014; Wang et al., 2018). We thus evaluated the correlation of the average timecourse of activation to each of our three tasks in both left and right IPS with the timecourse of each voxel of the brain. We note that the whole-brain comparison between the network functionally connected to the IPS during the arithmetic video compared to the grammar videos is reported in the Supplementary materials, while the following section only focuses on the two tasks that shared the same math content.

These ROI-to-voxel analyses first revealed that the network functionally connected to the IPS was larger during the math video lesson than during the controlled math task (math video: 91218 ± 128 voxels, math task: 81611 ± 112 voxels; paired t-test: t(31) = 2.40, p = 0.023). Moreover, in a whole-brain comparison of the naturalistic versus controlled math tasks, the IPS showed more connectivity with regions of the left temporal lobe and left inferior frontal gyrus, that are classically involved in language processing (Figure 5). Conversely during the controlled math task, the IPS appeared to be more correlated with a region of the right inferior frontal gyrus. Finally, we evaluated the connectivity strength of the IPS within the math-related network by averaging the correlation values extracted from the three remaining math-related regions of interest (bilateral pITG and right IFGOper). Converging with our previous results, a paired t-test revealed that the within math network connectivity was higher during the math video than during the math controlled tasks (math video: z = 0.40 ± 0.0069, math task: z = 0.37 ± 0.0079; paired t-test: t(31) = 3.72, p < 0.001).

**Figure 5:**
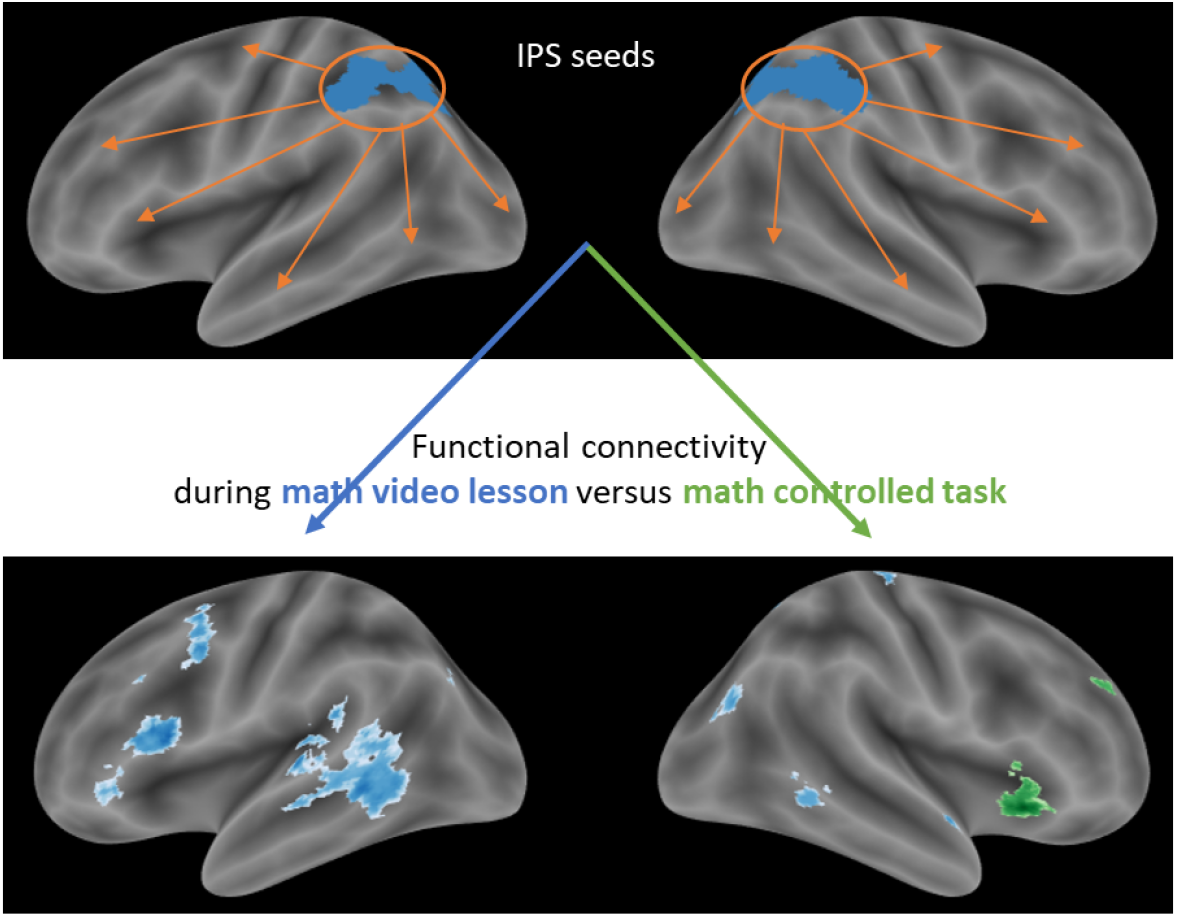
Brain regions functionally connected to the IPS during the naturalistic versus controlled math tasks. *Top*: Display of both left and right IPS regions that served as seeds in the ROI-to-voxel functional connectivity analysis. *Bottom*: Map of the brain regions exhibiting significant correlations with, i.e. functionally connected to, bilateral IPS, during the naturalistic math video lesson more than during the controlled math task (in blue). The reverse contrast is displayed in green on the same brain template. Both maps are displayed with p < 0.001 voxel-wise and FDR p < 0.05 cluster-wise.

These results suggest that the connectivity of the intraparietal sulcus was enhanced by the type of task (for the naturalistic more than the controlled math task), and converge with our analysis of entropy differences between tasks. These findings are thus compatible with the idea that long scale neural entropy might reflect increased connectivity between distributed brain regions.

### Increased neural entropy reflects advanced neural processing

We finally examined the question of whether the complexity of the neural signal increases with development. At the whole-brain level, we started by comparing the entropy of the neural signal elicited by our three tasks in adults versus children. Pulling all timescales together, we observed that adults’ neural responses generally had greater entropy than children’s throughout the brain for each task (top of figure S1). This result was confirmed by more precise analyses in our predetermined regions of interests. Figure S1 reveals that, particularly at short timescales, neural entropy was higher for adults compared to children in all regions of interest and for all tasks (all ps < 0.02). Thus, greater entropy is a signature of mature processing as opposed to disordered or immature processing.

To further assess this issue, we then examined the relation between children’s neural entropy and their neural maturity - that typically characterizes how “adult-like” is a child’s neural response. Neural maturity is commonly defined as the similarity between a child’s and the average adults’ neural responses, for each voxel of the brain and over the course of a given task (see Supplementary materials for a full analysis of neural maturity). During the math video lesson, at short timescales and to a lesser extent at long timescales, we observed that children’s neural entropy and neural maturity were significantly correlated in a large occipito-parietal network, including the bilateral IPS and pITG, and in the bilateral IFG Oper (Figure 6). We note here that these regions were remarkably similar to the math-related developing regions identified in 4 to 8-year-olds by Kersey et al. (2019) (see the overlap on Figure 6). Particularly at long timescales, we also observed significant correlations between neural entropy and neural maturity during the grammar video in the left IFG tri and STS, as well as at some occipito-parietal sites (see Figure S2).

**Figure 6:**
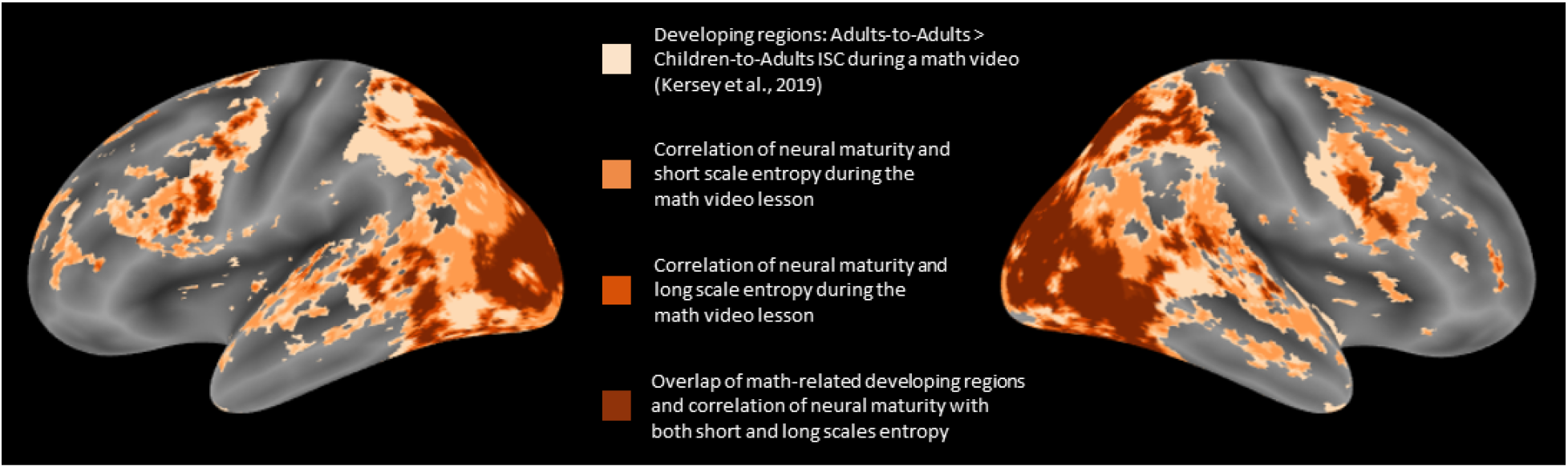
Correlation of neural entropy and neural maturity in developing regions. Overlap (in brown) of brain maps showing the developing math regions from the study by Kersey et al. 2019 (in light beige), and the correlation of neural maturity and entropy during the math video lesson (short scales in light orange and long scales in dark orange).

In our previously defined math-related ROIs, we finally compared the correlation between mean entropy and mean neural maturity values for each of our three tasks (Figure 7). At short timescales, during the math video lesson, the correlation was particularly strong in the right hemisphere (right IPS: Pearson’s coefficient r = 0.622, *R*^2^ = 0.387, p = 0.006; right pITG: r = 0.777, *R*^2^ = 0.604, p < 10-4; and right IFG Oper: r = 0.788, *R*^2^ = 0.621, p < 10-4), and slightly lower but still significant in the left hemisphere (left IPS: r = 0.487, *R*^2^ = 0.237, p = 0.040; and left pITG: r = 0.565, *R*^2^ = 0.319, p = 0.015). We note that children’s neural entropy and neural maturity were also significantly correlated in the right IPS during the controlled math task (r = 0.501, *R*^2^ = 0.251, p = 0.034). However, no math-related regions but the right IPS exhibited a correlation between neural entropy and neural maturity during the grammar video lesson (r = 0.525, *R*^2^ = 0.275, p = 0.025). We note that no such specificity was observed at long timescales (see Supplementary materials).

**Figure 7:**
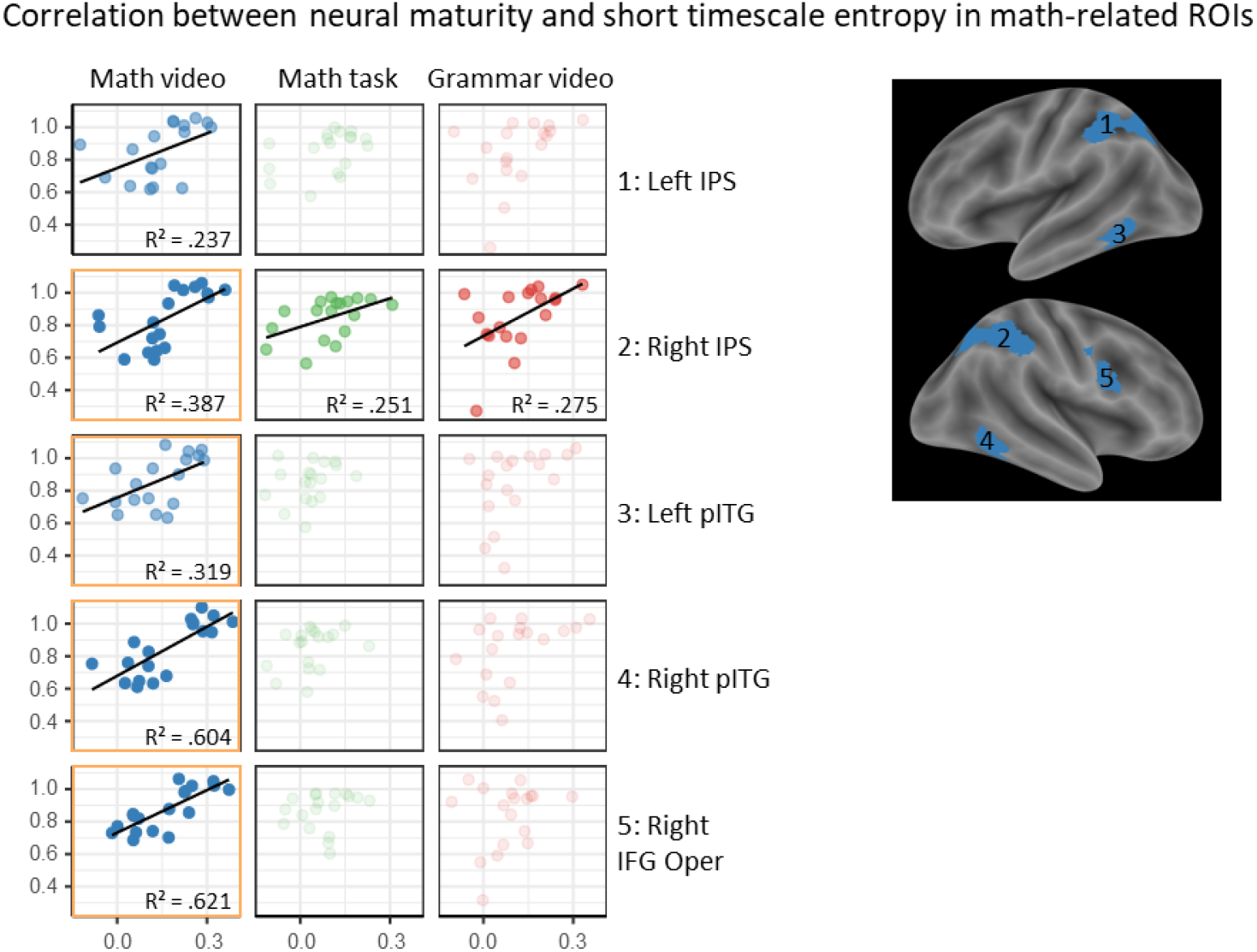
Neural maturity and neural entropy for math and grammar tasks. Evaluation of the correlation between neural maturity and short timescales (1 to 5) neural entropy during our three tasks in 5 math-related regions of interest that are represented on an inflated brain. The level of transparency of the graphs reflects the level of significance of the correlation: dark colors correspond to p < 0.01, medium colors correspond to p < 0.05, and light colors indicate that significance was not reached. Orange squares indicate correlations that were still significant after FDR correction for multiple comparisons.

These findings suggest that neural entropy increased with neural maturity in math-related regions. At short timescales, it was particularly the case for mathematics tasks, but not for the grammar video lesson. Together with the fact that neural entropy was generally greater in adults than children, our results are consistent with the hypothesis that increased entropy reflects advanced neural processing.

### Neural maturity is associated with decreased variance of the neural response

While neural entropy - that can be seen as a measure of temporal variability of the neural response - seems to increase over the course of development, is it also the case for the amplitude variability (i.e. variance) of the neural response? To answer this question, we computed the variance of the BOLD signal in each voxel of the brain, for each participant and during each task. A general ANOVA showed that children exhibited an overall greater variance than adults for the conjunction of both math tasks (Figure 8). This seems to indicate that, contrary to temporal complexity, the amplitude variability of neural responses tends to decrease with development.

**Figure 8:**
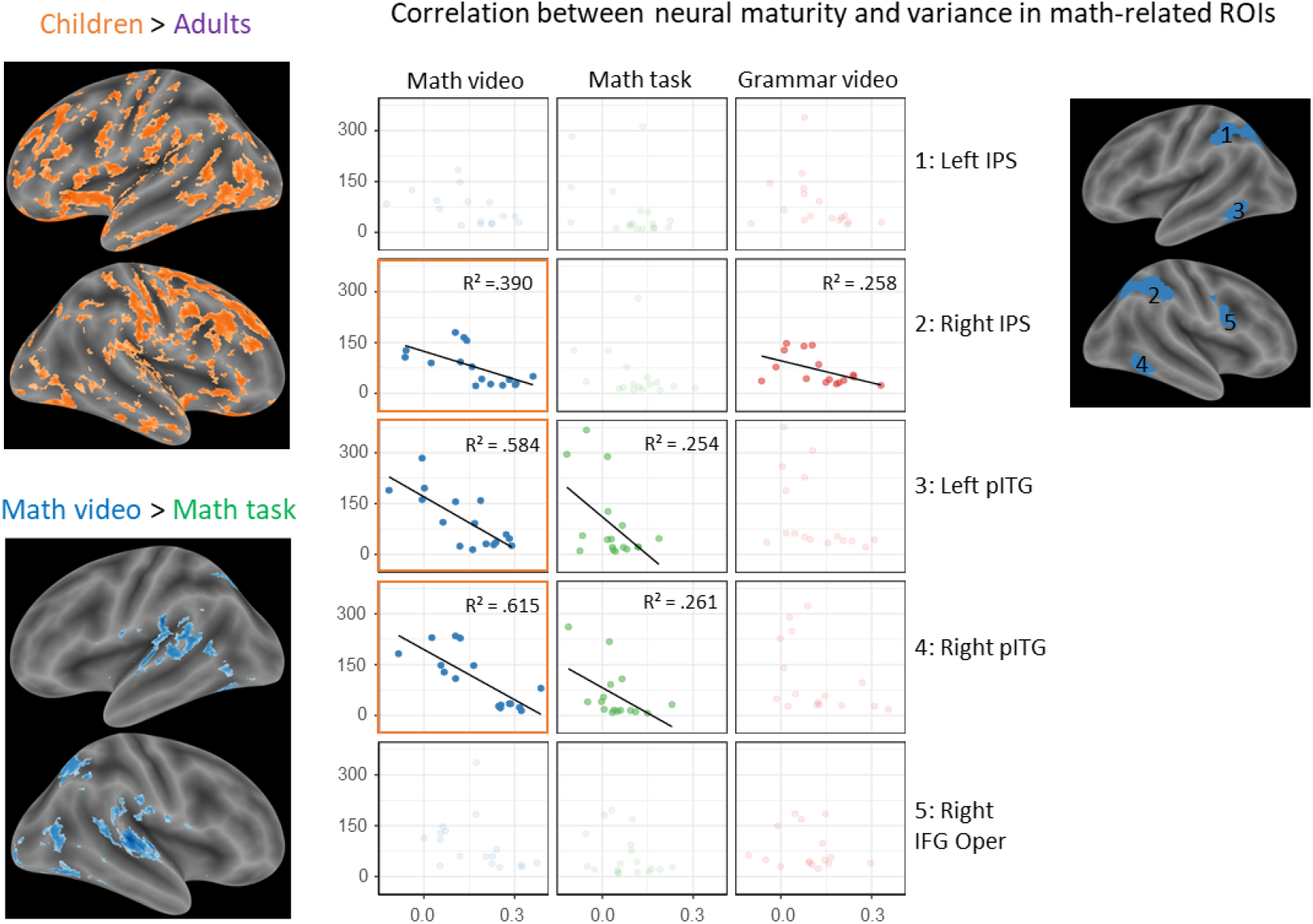
Neural maturity and variance for all tasks. Maps on the left show brain regions that exhibit higher amplitude variance for children compared to adults (top), and during the naturalistic math video compared to the controlled math task (bottom). Graphs on the right reveal the inverse correlation found between neural maturity and variance, in children, in our predefined math-related regions of interests. As in Figure 7, the level of transparency reflects the level of significance of the correlation: dark colors correspond to p < 0.01, medium colors correspond to p < 0.05, and light colors indicate that significance was not reached. Orange squares indicate correlations that were still significant after FDR correction for multiple comparisons.

We note in passing that the type of task also affected the variance of the neural response, but in a set of regions that differed greatly from what we observed so far. In fact, the naturalistic math task elicited greater variance than the controlled math task mostly in the primary visual and auditory cortices (in particular along the calcarine sulcus, and in bilateral Heschl gyri), and more generally along the superior temporal sulci of both hemispheres, in a large ventral occipital region, in the posterior cingulate gyrus, as well as in the precuneus.

As previously, we evaluated the correlation between mean variance and mean neural maturity values for each of our three tasks in our math-related ROIs (Figure 8). During the math video lesson, we found a significant inverse correlation in the right IPS (Pearson’s coefficient r = −0.625, *R*^2^ = 0.390, p = 9.69×10-3), the left pITG (r = −0.764, *R*^2^ = 0.584, p = 5.69×10-4), and the right pITG (r = −0.784, *R*^2^ = 0.615, p = 3.26×10-4; and right IFG Oper: r = 0.788, *R*^2^ = 0.621, p = 3.26×10-4). Children’s neural maturity was also significantly anti-correlated in both left and right pITG during the controlled math task (left: r = −0.504, *R*^2^ = 0.254, p = 0.039; right: r = −0.511, *R*^2^ = 0.261, p = 0.043). However, only the right IPS exhibited a correlation between variance and neural maturity during the grammar video lesson (r = −0.508, *R*^2^ = 0.258, p = 0.045). We note that the correlations found during the math video lesson were still significant after FDR correction for multiple comparisons. These findings altogether suggest that advanced neural processing goes with decreased variance of the neural response.

## Discussion

Natural learning requires integration of information over time. We compared children’s neural activity in a naturalistic education task in which mathematics learning is embedded in a building narrative versus neural activity during a more traditional two alternative forced choice task. Intersubject correlation measures revealed reliable neural signals across subjects in the IPS during the two types of mathematics tasks, and this pattern differed from that of the grammar task. Multiscale entropy is a measure that reveals changes in the complexity of neural signals over various timescales. Our data show that entropy was significantly greater at coarse timescales in the IPS during the naturalistic task compared to the laboratory task. This pattern was not observed for the naturalistic grammar task, suggesting that naturalistic mathematics tasks evoke neural integration of mathematics information over longer periods of time within the IPS. Furthermore, neural entropy in the parietal cortex increased with development and thus likely reflects advances in neural processing as opposed to disorder or noise.

Regions of the IPS have previously been implicated in semantic processing of mathematical concepts (e.g. Amalric and Dehaene, 2019, 2016; Cantlon and Li, 2013; Dehaene et al., 2003; Huth et al., 2016; Vogel et al., 2015; Zhang et al., 2012). For example, in 2012, Zhang et al. showed that the left IPS was involved in the semantic processing of geometric words, along with number words. More recently, studies by Amalric and Dehaene (2016, 2019) identified a network including bilateral IPS and pITG, that activated when participants processed spoken math statements and evaluated their truth value, in a systematic way that did not depend on the math domain, or statement difficulty. Moreover, in an fMRI study that systematically mapped semantic information contained in auditory narrative stories, Huth et al. (2016) revealed that bilateral parietal, inferior frontal and inferior temporal regions were particularly selective to words referring to numbers, units of measure, positions, and distances (see Figure 2 - Amalric and Dehaene, 2018).

Here we confirm and extend those prior findings by comparing two tasks in which the semantic content is the same but the format of the stimuli differs – a naturalistic educational lesson and a laboratory forced choice task on the same arithmetic principle. Both the naturalistic and laboratory mathematics tasks modulated neural activation in the IPS and dissociated from the non-mathematics grammar task. The brain networks for high-level math statements processing and simpler math computation were previously shown to overlap (Amalric and Dehaene, 2016), and to dissociate from the networks involved in the processing of equivalent non-math tasks (Amalric and Dehaene, 2018; Kersey et al., 2019; Monti et al., 2012). Thus, the naturalistic and laboratory mathematics tasks showed similarities in their underlying neural processes, and these similarities are likely semantic in nature.

Although there were similarities in neural processes between the naturalistic and laboratory mathematics tasks, there were also differences in their neural signatures. A key task difference between the naturalistic and controlled tasks is that in the naturalistic task, past information was needed to process incoming information over a longer period of time. The naturalistic task presented a virtual teacher explaining a mathematical concept over the course of several seconds whereas the laboratory task presented isolated stimuli in rigid task trials over a few seconds. In the naturalistic mathematics task, neural entropy was greater than entropy in the controlled mathematics task at long timescales. Prior studies have found that slow fluctuations in neural signals are sensitive to cumulative meaning. For example, Hasson et al. (2015) showed that higher-order temporal patterns of neural processing during naturalistic thought are hierarchical and functionally related to semantic input. They showed that slow fluctuations in neural signals depend on cumulative information processing over seconds to minutes and track the building meaning of a narrative over words, sentences, and events. Thus, the differences we observed in the neural entropy between the naturalistic and laboratory tasks within the IPS regions that were sensitive to the semantics of mathematics, likely reflect the accumulation of information over time promoted by the continuous educational narrative in the naturalistic task but not in the controlled task. Moreover, slow fluctuations in neural activity in the IPS were sensitive to the mathematics lesson more than to the grammar lesson. This indicates that the entropy in the neural signal within the parietal cortex was not the trivial reflection of naturalistic tasks complexity, but depended on stimulus content. Temporal entropy in a naturalistic educational task is thus a candidate marker of children’s semantic processing in the IPS.

Temporal entropy also appears to be a useful marker of development in children. As previously reported in the electrophysiology literature, the complexity of EEG signals increases over development during childhood (Miskovic et al. 2016), and from childhood to young adulthood (McIntosh et al., 2008). It has moreover been suggested that brain maturation is characterized by higher entropy of EEG signals at short time scales (Szostakiwskyj et al., 2017), and that it is accompanied by a long-scale entropy decrease as a potential consequence of a shift from global to local information processing (McIntosh et al., 2014; Wang et al., 2016). Here, for the first time in an fMRI study, we also observed that adults’ neural responses had greater entropy than children’s neural responses, especially at fine timescales.

Furthermore, in math-related regions such as IPS and pITG, that were similar to the math-related developing regions identified in 4 to 8-year-old by Kersey et al. (2019), we found that children’s neural entropy elicited by the math but not by the grammar task predicted their neural maturity. Finding such a correlation is far from trivial, as two signals with greater entropy do not necessarily become more correlated (see Figure S3 for examples of signals with dissociated correlation and entropy values). Arguably, such a correlation in the brain is even counter-intuitive. Indeed, one might expect that as one person’s neural signals become richer and more complex, they are less likely correlated with another person’s neural signals. However, our findings suggest that the complexity of neural signals not only increases over development but also increases in a systematic way, resulting in increased neural similarity among adults. In right IPS and pITG, increased neural complexity and similarity were also accompanied by decreased variance. Neural maturity, neural entropy and signal variance thus appear to be independent but complementary markers of brain development.

Lastly, our results altogether suggest that neural entropy reflects advanced neural processing. Converging evidence from electrophysiological studies reveal that enhanced neural entropy is associated with greater knowledge representations and information processing capacity (Beharelle et al., 2012; Heisz et al., 2012). Currently, neural entropy has been associated with the probability of neuronal firing within and between regions and their interconnectivity (Vakorin et al., 2011). Our findings that the IPS was functionally connected to a larger network during the naturalistic math task than during the controlled math task are compatible with this idea. Previous ECoG, EEG, and fMRI studies with adults have shown that the entropy of neural activity within a region scales with how interconnected that region is to other regions of the brain (Wang et al., 2018). Larger and more interconnected networks produce greater variation in neural responses over time, or entropy. Increased entropy may thus reflect a wider repertoire of brain states within or between network, or greater functional distribution of those states across networks (Bassett and Bullmore, 2006; Deco et al., 2011; Freeman, 2000). In light of these prior findings, the correlation we observed between children’s neural entropy and neural maturity within the math network may reflect the increasing breadth and diversity of neural processes available to children for analyzing mathematical information over development.

## Methods and Materials

### Participants

38 subjects participated in the study. 24 typically developing children at the end of their 3rd grade year and 14 adults participated in this study. 6 children were excluded from further analyses because of excessive head-motion or opting-out.

### Ethics statement

All participants gave informed consent after reading or being read consent information. The protocol was approved by both the University of Rochester and the Carnegie Mellon Institutional Review Boards.

### Procedure

Each trial of the same/different task started with two white circles horizontally distributed on a gray background, and a blue fixation cross at the center of the screen. After 500 ms, the fixation cross disappeared, and a first operation or dot array appeared in the first circle. At 1500 ms, a second operation or dot array appeared in the second circle. 1 second later, the two circles turned to red, indicating the beginning of the 4 seconds response period. The two operations or dot arrays remained on the screen during the entire response period. Participants were instructed to press a button with their right thumb if they thought that the two operations had the same result or the two arrays had the same number of dots, and to press a button with their left thumb otherwise. Each trial ended with a rest period of variable duration (mean = 5s, sd = 1s).

Both the first and second session of the test included 3 runs of 24 trials each. In order to perform inter-subject correlations, the order of trials was fixed within a run, but we randomized the order of runs.

Movie runs started with a plain blue screen with a central black fixation cross for 10 seconds, in order to measure the activation baseline before the movie started playing. Both movies consisted of three video sequences of 80 ± 4 seconds, separated by 10 seconds rest periods. No specific instruction was given to participants except watching the movie. Half participants watched the math movie first and the other half watched the grammar movie first.

### Stimuli

#### Pedagogical videos

We created two videos with the online application “Powtoon”. The videos respectively explained the commutative principle of multiplication and relative pronouns. Both movies were visually matching (same blue background, same cartoon-like animations, same amount of information on screen, same combination of symbols and pictures).

#### Numerical pairs

The same pairs were presented symbolically and non-symbolically in each run. Symbolic pairs were simple math operations (e.g. “2×3” and “3×2”). Their non-symbolic counterparts were sets of dots visually arranged in subgroups (e.g. ‘2 groups of 3 dots’ and ‘3 groups of 2 dots’). Subgroups of the same size in a set were arranged according to the same pattern. In each run, 8 pairs tested for the understanding of the commutative principle (e.g. “4×2” and “2×4”). We compared them to 8 pairs with the same number of items but different math operations (e.g. “4+2” and “3×2”). We also introduced 8 pairs controlling for numerosity and ensuring that subjects did not simply estimate the number of items but computed exact additions and multiplications (half of them used the same operation: e.g. “4×3” and “4×2”; and the other half used different operations: e.g. “5+2” and “2×3”). See Table S1 for the full list of stimuli.

#### fMRI procedures

Functional images were acquired on a Siemens Prisma 3-Tesla scanner at the Rochester Center for Brain Imaging with multiband imaging sequences (multiband factor = 4, slice acceleration factor = 3, 60 interleaved axial slices, 3 mm thickness and 3 mm in-plane resolution, TR = 2000 ms, TE = 30 ms), with 64 channel head-coil.

### Analyses of fMRI data

#### Preprocessing

fMRI data were processed with SPM12. Functional images were first corrected for slice timing, realigned, normalized to the standard MNI brain space, resampled to a 2 mm voxel size, and spatially smoothed with an isotropic Gaussian filter of 4mm FMWH. The 6 framewise displacement parameters extracted from previous realignment were regressed out, and a 100-second high-pass filter was applied. The relevant segments of each subject’s data were then trimmed (to remove the first seconds of each epoch capturing nonspecific stimulus onset) and z-scored (standardized to zero mean and unit variance) for each voxel and segment. They were finally concatenated into three 4D NIfTI files: arithmetic video lesson, grammar video lesson, and math forced-choice task. An average gray-matter mask was applied prior to performing any further analyses.

#### Intersubject correlations

The technique of intersubject correlation (ISC) is a data-driven analysis technique which consists in identifying brain regions where the response to a stimulus is systematic over time, i.e. correlated among participants (Nastase et al., 2019). Both within groups (children and adults) and between children and adults, the intersubject-correlation was calculated using the leave-one-out approach. At each voxel, each subject’s time series was correlated with the average time series of all other subjects. This approach directly gave us one R-map for each subject, that was converted into a z-map using the Fischer transformation. Note that the correlation between each child and the group of adults is typically interpreted and referred to as the child’s neural maturity. At the group level, we then performed t-tests and ANOVAs on the z-maps of each task. When pulling all participants together, we performed t-tests and ANOVAs on adults-to-adults and children-to-adults correlation maps.

#### Multiscale entropy of BOLD signal

The multiscale entropy (MSE) was proposed to estimate the dynamic complexity in a time series by considering different time scales. The MSE of the signal for the three tasks was calculated for each participant at each voxel included in the gray matter mask. To do so, we used the WFDB toolbox (https://physionet.org/physiotools/mse/) that implements a two-steps analysis described by Costa et al. (2002). In this method, multiple “coarse-grained” time series of fixed length L/n are first formed by averaging n consecutive non-overlapping data points of the initial time series of length L (see Figure 1). Then the sample entropy of each coarse-grained time series is calculated. The sample entropy (SampEn) is an approximation of Kolmogorov complexity that has been proposed for physiological signals. It essentially consists in determining how reproducible is a pattern of length m throughout the entire time series. It proceeds first by counting how many times each pattern, of length m and m+1 respectively, repeats across the time series, considering a tolerance factor r. In more details, two patterns [x1,x2] and [y1,y2] are considered equal if max(|xi-yi|) < r (see Figure 1, bottom panel). Second, SampEn is calculated as the negative natural logarithm of the ratio between these two counts. The different parameters were chosen according to the optimal parameters found in a previous fMRI study of adolescents (Easson and McIntosh, 2019): pattern length m = 2; tolerance factor r = 0.5; scale factor of coarse-graining l = 10. This gave us 10 E-maps (one for each scale) for each task and each participant. These E-maps were then averaged over short scales (1 to 5) and long scales (6 to 10) and entered in a repeated measure ANOVA to evaluate group-level whole-brain maps. Note that unless otherwise specified, maps are displayed at p < 0.001 voxelwise and p < 0.05 clusterwise, FDR corrected.

#### Analysis of variance of the neural signal

To compute the variance of the neural signal, we used preprocessed data centered to the mean but not normalized to unit variance. At each voxel and for each task (arithmetic video lesson, grammar video lesson, and concatenated math forced-choice task), each subject’s time series was extracted and its variance computed. At the group level, we then performed an ANOVA on the resulting maps of variance.

#### ROI analyses

Math- and language-related regions were defined from two completely independent studies. From the study of the constituent structure of sentences by Pallier et al. (2011), we selected 4 (out of 6) left-hemispheric ROIs: the anterior superior temporal sulcus (aSTS), the temporo-parietal junction (TPJ), and the inferior frontal gyrus pars orbitalis (IFG Orb) and pars triangularis (ITF Tri). We note here that we chose left-hemispheric aSTS and TPJ over the temporal pole (TP) and posterior superior temporal sulcus (pSTS) because bilateral aSTS and TPJ are also typically involved in semantic processing (Binder et al., 2009). The math-related regions were the ones previously used by Amalric and Dehaene (2016) and identified as processing even advanced math concepts. In more details, 5 regions in the bilateral intraparietal sulci (IPS), bilateral posterior inferior temporal gyri (pITG), and the right inferior frontal gyrus pars opercularis (IFG Oper) were defined from the contrast of calculation versus sentences of localizer scans performed in a cohort of 83 subjects (Pinel et al., 2007).

Each of the 9 above regions were then flipped to the other hemisphere, thus defining 4 language right-hemispheric symmetrical regions in aSTS, TPJ, IFG Orb, and IFG tri, and 1 math left-hemispheric symmetrical region in IFG Oper. Finally IPS and pITG regions were defined as the common voxels between the original region and its flipped counterpart. This construction thus resulted in 14 symmetrically paired regions of interest in which we performed various types of analyses.

Within each region and for each task, we first extracted the intersubject correlation values for each adult (relative to the other adults), and for each child (both relative to the other children and relative to the adults). For each adult and children participant, we also extracted the entropy values for each scale. These values were either averaged and entered into traditional t-tests, or directly used for further analyses.

For the lateralization analysis, we considered a combination of both the intensity and extent of intersubject correlations, by evaluating the sum of positive ISC values (∑ISC+) over each region and for each participant. For each pair of regions, ISC lateralization indices were then calculated following the classical formula LI = (∑ISC+lh - ∑ISC+rh)/(∑ISC+lh + ∑ISC+rh). LI values greater than 0.1 typically characterize a left-lateralization while LI values less than −0.1 typically characterize a right-lateralization. Values close to 0 are characteristic of a bilateral distribution.

Representational similarity analyses were performed only in the math-related regions of interest (left and right IPS, left and right pITG, and right IFG Oper). This multi-voxel pattern analysis typically allows to measure the spatial similarity of the BOLD signal elicited by various stimuli. Here, we applied it to test how spatially similar was the synchronous neural activity elicited by our three tasks. To do so, we evaluated for each participant the correlation of ISC values during our three tasks over all the voxels of each math-related ROI. We then used t-tests to compare the correlation values of both math tasks versus the correlation values of both video lessons.

To test the correlation between children’s neural maturity and neural complexity, we first averaged the mean entropy values over short timescales (scales 1-5) for each child and for each primarily defined regions of interest (i.e. left and right IPS, left and right pITG, right IFG Oper, left aSTS, left TPJ, left IFG Orb and left IFG tri) to which we added the anatomically defined left fusiform gyrus. We then evaluated the correlation between these entropy values and the mean intersubject correlation values for each child relative to the group of adults.

Note that a Bonferroni correction or FDR correction for the number of ROIs (or pairs of ROIs) was applied to all test results.

#### Functional connectivity analysis

Preprocessed BOLD data were analyzed with the CONN functional connectivity toolbox (Whitfield-Gabrieli and Nieto-Castanon, 2012). Both left and right IPS (defined above) were used as seeds in a whole-brain analysis. In this seed-to-voxel analysis, the toolbox evaluates the Fisher-transformed bivariate correlation values between the BOLD timeseries averaged across the seed ROI and the BOLD timeseries of each individual voxel.

## Acknowledgments

The authors would like to thank Alyssa Kersey, Lorenzo Ciccione, and Randy McIntosh for sharing respectively their results, stimuli and methods.

