## Supplementary materials for "Entropy, complexity, and maturity in children’s neural responses during naturalistic mathematics learning"

***Complementary behavioral results***

Excluding missed trials, global ANOVAs revealed in children and adults alike, a main effect of the pair categories on accuracy, and main effects of both symbolism and the pair categories on reaction time (all p*s* < 0.003). In particular, numerically identical pairs involving different math operations induced more errors than pairs involving the same math operation either numerically identical (children: t(17) = 4.11; adults: t(13) = 4.34; both p*s* < 0.001) or not (children t(17) = 3.35; adults: t(13) = 2.90; both p*s* < 0.02), and also elicited slower responses (children: t(17) = 8.71 and 2.84 respectively; adults: t(13) = 8.06 and 2.19 respectively; all ps < 0.05). In addition, children and adults consistently answered to symbolic trials quicker than to non-symbolic trials (children: t(17) = 8.82; adults: t(13) = 5.41; both p*s* < 0.0005).

After watching the first video lesson (either math or grammar) and before watching the other video lesson, all participants were tested again on the same/different task. In this second test, children who just watched the arithmetic video answered slightly but not significantly faster than children who watched the grammar video (arithmetic video group: 1.77 ± 0.101s, grammar video group: 1.92 ± 0.125s, t(16) = 0.96, p = 0.35). A similar effect was observed in adults (arithmetic video group: 0.864 ± 0.125s, grammar video group: 1.07 ± 0.195s, t(12) = 0.98, p = 0.34). No significant difference between groups was found on accuracy neither in children nor in adults. We then analyzed the evolution of performance between the first and second tests for each category of pairs. To do so, we calculated the differences of accuracies and ratios of reaction times for each category and each subject. In both children and adults, ANOVAs with symbolism and pair categories as within factors, the first video watched as between factor, and participants as a random effect did not reveal any significant improvement of accuracy or reaction time for the group who watched the arithmetic movie first. However, a significant accuracy increase and a reaction time decrease were found only in the group of children who watched the arithmetic video first, for non-symbolic pairs that could be seen as exemplars of the commutative principle, i.e. numerically equivalent and both involving multiplication (accuracy: t(8) = 2.58, p = 0.016; and reaction time: t(8) = 2.66, p = 0.014).

***Lateralization analysis***

For each child and adult participant, in 4 ROIs associated with language processing (aSTS, TPJ, IFG tri, and IFG Orb, from Pallier et al., 2011), we computed the lateralization indices (LI) observed during the grammar video. Particularly in frontal regions, while adults were significantly left-lateralized (IFG Orb

LI = 0.32 ± 0.056, t-test of LI > 0.1: $t(13) = 4.04$, $p < 0.001$; IFG tri LI = 0.45 ± 0.055, $t(13) = 6.63$, $p < 0.001$), children exhibited bilateral responses (IFG Orb LI = 0.091 ± 0.10, $t(17) = 0.088$, $p = 0.53$; IFG tri LI = 0.097 ± 0.064, $t(17) = 0.047$, $p = 0.52$). A between-group t-test confirmed that adults' neural responses to the grammar video were more left-lateralized than children's (IFG Orb: $t(30) = 1.86$, $p = 0.036$; IFG tri: $t(30) = 4.18$, $p < 0.001$). No such differences were found in TPJ and aSTS where both children and adults showed bilateral synchronous neural responses (all |LI*s*| < 0.21, all p*s* > 0.056). In aSTS, adults' neural responses were even more bilateral than children's (two-sample t-test: $t(30) = 2.85$, $p = 0.004$).

We then extended this lateralization analysis to neural responses elicited by the arithmetic video using 3 ROIs classically associated with math processing (IPS, pITG, IFG Oper) that were previously used by Amalric & Dehaene (2016) and defined from the study by Pinel et al. (2007). We observed bilateral responses in children and adults alike (all |LI*s*| < 0.074, all p*s* > 0.68, and two-sample t-tests were not significant).

### *Brief description of all ISC overlaps between tasks*

First, all three tasks included mostly visual inputs and exhibited a strong overlap (light green areas on Figure 3) of neural response in the occipital cortex, fusiform gyri, precuneus and inferior parietal lobule. In adults, this general overlap across all tasks also extended to the left inferior frontal gyrus pars opercularis.

The arithmetic and grammar pedagogical videos had both visual and auditory inputs and overlapped (pink areas on Figure 3) in the occipital cortex, lingual gyri, bilateral superior and middle temporal gyri, and bilateral inferior frontal sites (particularly in the left hemisphere in adults).

The naturalistic and controlled math tasks with common content overlapped (light blue areas on Figure 3) alongside the bilateral intraparietal sulcus (particularly in the right hemisphere), and in middle frontal regions in children and adults. In adults, we observed overlapping intersubject correlations also in bilateral posterior inferior temporal regions and in the mesial occipital cortex.

Finally, we observed some overlap between neural responses elicited by the controlled math task and the naturalistic grammar video (orange areas on Figure 3), at various frontal sites in adults and children, and at the temporo-parietal junction close to the angular gyrus in both hemispheres in children.

***Detailed analysis of correlations of neural responses between children and adults.***

We started by analyzing the neural correlation between children and adults (i.e. children's adult-like neural responses, often interpreted as children's neural maturity) in the contrast of arithmetic versus grammar video lessons. We observed right-lateralized mature responses at an inferior temporal site, in the parietal cortex, and in the superior frontal gyrus. The converse contrast of grammar versus arithmetic video lessons yielded correlations between children and adults in small clusters at the right temporo-parietal junction, and in the right middle frontal gyrus, as well as in the left IFG pars triangularis.

Then, we evaluated the overlap of correlated neural responses between children and adults for both math tasks. Similarly to what was observed among children and adults separately, both types of math tasks elicited remarkably overlapping neural responses in the bilateral intraparietal sulci, especially in the right hemisphere. Noticeably, while intersubject correlations among children alone did not reveal much synchronized neural responses in the posterior inferior temporal gyrus, children's neural responses in a right inferior temporal site appeared very similar to adults during math tasks.

Children's adults-like neural responses that were common to both arithmetic and grammar video lessons were found mostly in the occipital cortex, lingual gyri, bilateral superior and middle temporal gyri, and bilateral inferior frontal sites. This suggests that visual and auditory processing of the videos happened similarly in children and adults. Though in this case, the general overlap of all three tasks was minimally found in the inferior and middle occipital cortex, the fusiform gyri, and in two bilateral small sites of the posterior inferior parietal lobule. We also note that, contrary to what was observed among children and adults separately, the controlled math task did not yield much correlated activation between children and adults in the motor cortex. This might be due to the fact that children answered significantly slower than adults, thus introducing temporal differences between both groups in the motor cortex activity.

Furthermore, we evaluated which brain regions were showing differential intersubject correlations between children and adults. We first noted that adults systematically exhibited greater intersubject correlations among themselves than children among themselves. A subject-level ROI analysis showed that adults' ISC was significantly higher than children's during the controlled math test in both left and right IPS (p*s* < 0.0008, which is below the significance threshold of p = 0.0055 after Bonferroni correction for multiple comparisons in 9 ROIs). It was also the case during the arithmetic video lesson, in both left and right pITG (p*s* < 0.0001), and during the grammar video lesson in all left lateralized language ROIs (p*s* < $10^{-5}$ except in TPJ where p = 0.008). These results were confirmed at whole-brain level, as displayed on the top row of the figure below.

We finally sought to identify developing regions of the brain which, as suggested by Kersey et al. (2019), can be done by testing for greater intersubject correlations among adults than between children and adults. Noticeably, whole-brain t-tests for our three tasks (bottom row) revealed virtually the same regions as the ones identified by the direct comparison between intersubject correlations among adults and intersubject correlations among children (top row). During the arithmetic video lesson, adults exhibited stronger correlations among themselves than with children in a left inferior temporal region, bilaterally along the anterior superior temporal sulcus, in the right parietal cortex, at the left temporo-parietal junction. During the grammar video lesson, we observed mostly left-lateralized regions, including the superior temporal sulcus, the inferior parietal lobule, the fusiform gyrus and an extended swath in the inferior frontal gyrus. Finally, during the controlled math task, adults' neural responses were more correlated among themselves than with children bilaterally in the intraparietal sulcus, the motor cortex, the inferior frontal gyrus pars opercularis and the middle occipital cortex. These results were confirmed by t-tests performed on mean ISC values in our five math-related and four language-related predefined regions of interest (math: left and right IPS, left and right pITG, right IFG Oper; language: left aSTS, TPJ, IFG Orb, IFG tri).

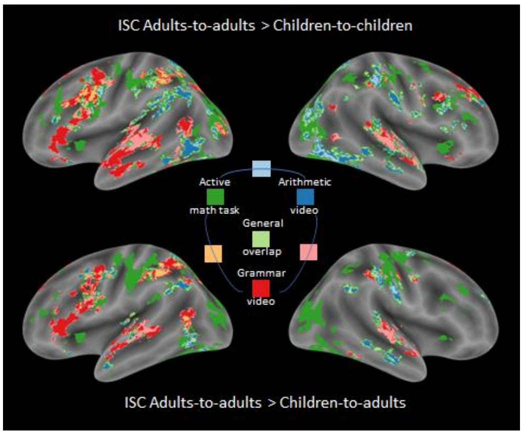


***Comparison of IPS functional connectivity for both educational videos***

The math video lesson revealed quite different patterns of functional connectivity than the grammar video lesson. At the whole-brain level, the IPS showed higher connectivity with the occipital cortex and an extended right inferior temporal region during the math video. During the grammar video, the IPS showed higher connectivity with bilateral aMTG and angular gyri, classically associated with semantic processing (Binder et al., 2009). Though we note that neither the overall extent of the network connected to the IPS, nor the global connectivity strength differed between both video lessons.

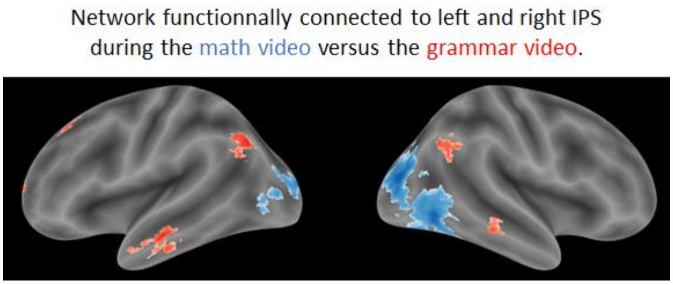

***Correlation between neural maturity and neural entropy in language-related ROIs***

At both short and long timescales, most language-related regions exhibited a significant correlation between neural entropy and neural maturity during the grammar video lesson (left fusiform gyrus: $r = 0.614$, $R^2 = 0.377$, $p = 0.007$; left TPJ: $r = 0.533$, $R^2 = 0.284$, $p = 0.023$; and the left aSTS: $r = 0.579$, $R^2 = 0.335$, $p = 0.012$). At short timescales, but not at long timescales, we also found correlations between neural entropy and neural maturity during the math tasks in some language-related regions such as the left TPJ, aSTS and IFG tri for the math video lesson ($0.479 < rs < 0.568$, $0.229 < R^2s < 0.322$, $0.05 > ps > 0.02$) or the left fusiform gyrus for the controlled math test ($r = 0.593$, $R^2 = 0.352$, $p = 0.009$).

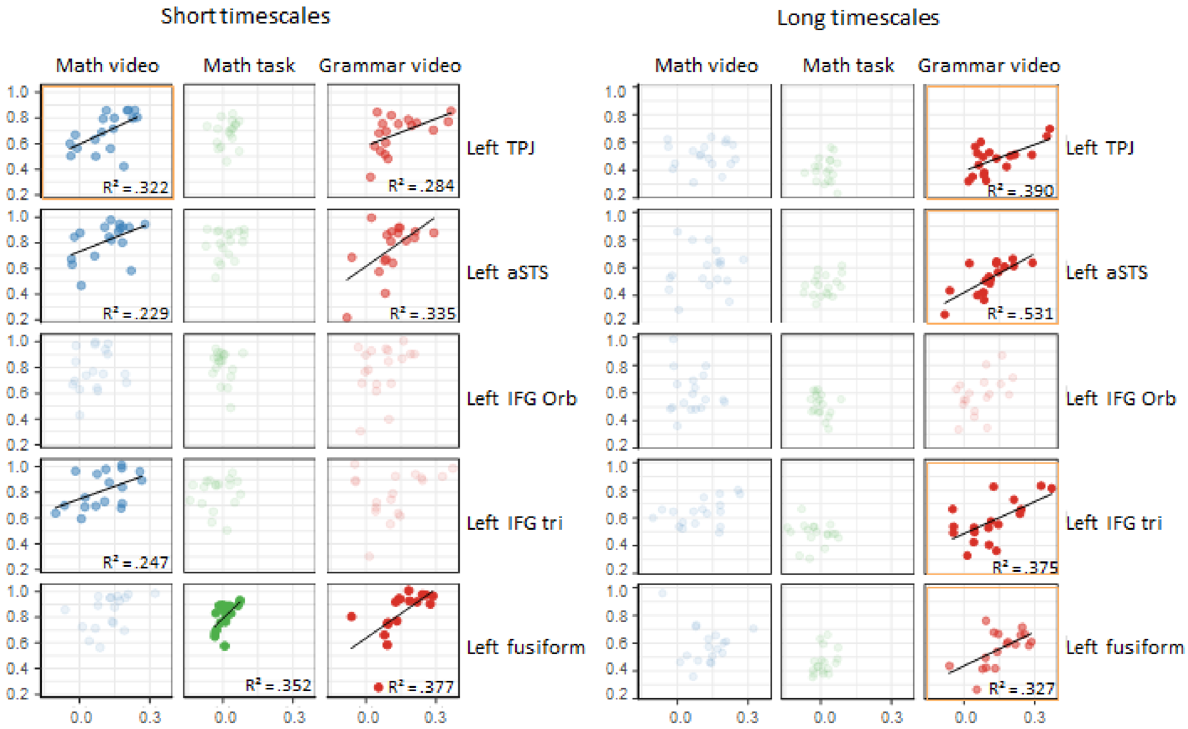

***Table S1: Pairs of sets and operations used in the forced-choice math task***

| # | 2nd item | 1st item | | | |
|---|---|---|---|---|---|
| | | Same number - Different set number | | Different number - Same set number | |
| | | Same operation | Different operation | Same operation | Different operation |
| 6 | 2x3 | 3x2 | 1+2+3 | 2x4 | 5+2 |
| | 3x2 | 2x3 | 4+2 | 3x4 | 3+4+5 |
| 8 | 2x4 | 4x2 | 1+2+2+3 | 2x2 | 1+3 |
| | 4x2 | 2x4 | 1+3+4 | 4x3 | 2+3+3+4 |

***Table S2. Brain regions exhibiting higher long scale entropy for the naturalistic math task than for the controlled math task.***

Table shows all local maxima separated by more than 20 mm, for t > 3.1455; p < 0.0010; df = 150; minimum extent = 35. Regions were automatically labeled using the AAL2 atlas. x, y, and z are the Montreal Neurological Institute (MNI) coordinates in the left-right, anterior-posterior, and inferior-superior dimensions, respectively.

| Adults | | | | | | Children | | | | | |
|---|---|---|---|---|---|---|---|---|---|---|---|
| Region Label | Extent | t-value | x | y | z | Region Label | Extent | t-value | x | y | z |
| Occipital_Inf_R | 45022 | 8.190 | 34 | -82 | -4 | Calcarine_R | 17976 | 7.858 | 12 | -92 | 4 |
| Occipital_Mid_L | 45022 | 7.776 | -34 | -92 | 2 | Occipital_Mid_L | 17976 | 7.285 | -32 | -94 | -6 |
| Lingual_L | 45022 | 7.728 | -10 | -84 | -10 | Occipital_Inf_R | 17976 | 6.122 | 38 | -68 | -10 |
| Temporal_Inf_R | 319 | 5.079 | 44 | 2 | -18 | Precentral_R | 812 | 5.733 | 46 | 4 | 28 |
| Temporal_Inf_R | 319 | 4.224 | 46 | 6 | -40 | Frontal_Inf_Tri_R | 812 | 5.174 | 52 | 22 | 20 |
| Cerebelum_7b_L | 400 | 5.024 | -24 | -70 | -46 | Postcentral_R | 4205 | 5.428 | 44 | -38 | 62 |

| | | | | | | | | | | | |
|---|---|---|---|---|---|---|---|---|---|---|---|
| Frontal_Mid_2_L | 173 | 4.764 | -46 | 42 | 16 | Temporal_Sup_R | 4205 | 5.216 | 68 | -24 | 10 |
| Frontal_Mid_2_L | 173 | 3.932 | -46 | 46 | -10 | Rolandic_Oper_R | 4205 | 5.080 | 46 | -26 | 22 |
| Frontal_Med_Orb_R | 193 | 4.758 | 6 | 50 | -12 | Cerebelum_8_R | 485 | 5.155 | 32 | -68 | -50 |
| Rectus_L | 193 | 4.163 | -2 | 30 | -20 | Cerebelum_Crus2_R | 485 | 4.536 | 22 | -80 | -36 |
| Rectus_L | 67 | 4.570 | -8 | 22 | -26 | Thalamus_R | 50 | 5.135 | 16 | -32 | 2 |
| Cerebelum_8_R | 159 | 4.475 | 32 | -70 | -56 | Supp_Motor_Area_L | 478 | 5.109 | -4 | -8 | 54 |
| Frontal_Sup_2_L | 51 | 4.418 | -24 | 62 | 0 | Supp_Motor_Area_L | 478 | 3.783 | -6 | 14 | 58 |
| OFCmed_R | 84 | 4.297 | 14 | 26 | -20 | Precentral_R | 175 | 5.002 | 28 | -16 | 62 |
| Insula_L | 47 | 4.186 | -36 | 14 | 6 | Frontal_Sup_2_R | 175 | 3.504 | 26 | 4 | 56 |
| Cerebelum_Crus1_R | 45 | 4.167 | 20 | -74 | -34 | Cerebelum_7b_L | 260 | 4.801 | -18 | -72 | -46 |
| Temporal_Pole_Mid_L | 45 | 4.065 | -32 | 10 | -34 | Cerebelum_Crus2_L | 260 | 4.134 | -38 | -66 | -46 |
| ParaHippocampal_R | 47 | 4.015 | 24 | 2 | -36 | Thalamus_L | 111 | 4.763 | -10 | -32 | 2 |
| Thalamus_L | 60 | 3.975 | -16 | -32 | 0 | Insula_R | 60 | 4.736 | 42 | 6 | -10 |
| Temporal_Inf_L | 84 | 3.946 | -54 | -22 | -22 | Thalamus_L | 37 | 4.449 | -8 | -18 | 14 |
| Rectus_L | 82 | 3.916 | -6 | 44 | -22 | Cingulate_Mid_L | 40 | 4.282 | 0 | 22 | 32 |
| Temporal_Pole_Sup_L | 53 | 3.795 | -36 | 6 | -20 | Temporal_Mid_L | 43 | 4.130 | -58 | -2 | -24 |
| Frontal_Mid_2_R | 45 | 3.776 | 44 | 34 | 28 | Cerebelum_Crus2_L | 53 | 3.962 | -14 | -76 | -36 |
| Temporal_Pole_Sup_L | 37 | 3.773 | -54 | 8 | -16 | Insula_R | 62 | 3.882 | 40 | 0 | 12 |
| | | | | | | Angular_R | 50 | 3.830 | 40 | -56 | 24 |
| | | | | | | Rolandic_Oper_R | 40 | 3.689 | 58 | 8 | 10 |

***Table S3. Brain regions exhibiting higher long scale entropy for the arithmetic video lesson than for the grammar video lesson.***

Same legend as Table S2.

| Adults | | | | | | Children | | | | | |
|---|---|---|---|---|---|---|---|---|---|---|---|
| Region Label | Extent | t-value | x | y | z | Region Label | Extent | t-value | x | y | z |
| Parietal_Sup_L | 5468 | 6.873 | -20 | -66 | 60 | Occipital_Mid_L | 2102 | 7.122 | -32 | -94 | -6 |
| Postcentral_R | 5468 | 6.211 | 46 | -32 | 54 | Fusiform_L | 2102 | 6.064 | -40 | -64 | -12 |
| Occipital_Mid_L | 5468 | 5.956 | -28 | -68 | 34 | Occipital_Mid_L | 2102 | 5.446 | -24 | -90 | 16 |
| Precentral_L | 373 | 6.167 | -52 | 8 | 32 | Occipital_Inf_R | 1552 | 6.672 | 28 | -96 | -4 |
| Calcarine_L | 294 | 5.838 | 0 | -94 | 0 | Occipital_Inf_R | 1552 | 5.966 | 42 | -68 | -10 |
| Lingual_L | 294 | 5.127 | -14 | -94 | -16 | Lingual_R | 1552 | 5.179 | 18 | -78 | -6 |
| Cerebelum_Crus1_L | 294 | 4.880 | -40 | -70 | -22 | Temporal_Sup_L | 1867 | 5.767 | -62 | -22 | 6 |
| Temporal_Inf_R | 233 | 5.739 | 54 | -52 | -8 | Temporal_Mid_L | 1867 | 5.245 | -56 | -48 | 14 |
| Temporal_Sup_L | 616 | 5.558 | -52 | -16 | 6 | Postcentral_L | 1867 | 5.231 | -48 | -24 | 52 |
| Rolandic_Oper_L | 616 | 4.555 | -40 | -34 | 14 | Precentral_R | 258 | 5.733 | 46 | 4 | 28 |
| SupraMarginal_L | 139 | 5.459 | -60 | -24 | 40 | Frontal_Mid_2_R | 258 | 3.691 | 38 | 24 | 22 |
| Temporal_Sup_R | 755 | 5.297 | 50 | -28 | 10 | Fusiform_L | 31 | 5.520 | -28 | -36 | -24 |
| Temporal_Sup_R | 755 | 4.318 | 64 | -12 | 4 | Postcentral_R | 584 | 5.428 | 44 | -38 | 62 |

| | | | | | | | | | | | |
|---|---|---|---|---|---|---|---|---|---|---|---|
| Frontal_Inf_Tri_L | 93 | 5.267 | -50 | 16 | 6 | Postcentral_R | 584 | 4.176 | 52 | -20 | 54 |
| Temporal_Sup_R | 38 | 5.224 | 64 | -50 | 20 | SupraMarginal_R | 584 | 4.017 | 38 | -34 | 42 |
| Temporal_Inf_L | 80 | 5.170 | -52 | -58 | -18 | Temporal_Sup_R | 440 | 5.214 | 68 | -24 | 10 |
| Lingual_L | 205 | 5.143 | -16 | -48 | -2 | Temporal_Sup_R | 440 | 4.305 | 52 | -46 | 16 |
| Calcarine_L | 205 | 3.992 | -18 | -66 | 10 | Temporal_Sup_R | 440 | 3.885 | 58 | -6 | 0 |
| Frontal_Mid_2_L | 112 | 5.114 | -28 | 10 | 64 | Frontal_Inf_Tri_R | 175 | 5.170 | 52 | 22 | 20 |
| Cerebelum_7b_L | 96 | 5.007 | -24 | -70 | -46 | Cerebelum_8_R | 224 | 5.138 | 32 | -68 | -50 |
| Precentral_R | 60 | 5.005 | 30 | -24 | 64 | Cerebelum_Crus2_R | 224 | 4.093 | 22 | -82 | -36 |
| Temporal_Mid_L | 52 | 4.907 | -68 | -40 | -2 | Frontal_Inf_Tri_L | 885 | 5.100 | -46 | 18 | 30 |
| Calcarine_L | 17 | 4.815 | -12 | -82 | 10 | Precentral_L | 885 | 4.595 | -38 | -4 | 46 |
| Cuneus_L | 95 | 4.796 | -16 | -70 | 22 | Precentral_L | 107 | 5.077 | -32 | -12 | 54 |
| Precentral_R | 199 | 4.765 | 56 | 6 | 36 | Rolandic_Oper_R | 279 | 5.075 | 46 | -26 | 22 |
| Precentral_R | 199 | 4.396 | 40 | -6 | 48 | SupraMarginal_R | 279 | 4.801 | 66 | -18 | 30 |
| Cuneus_L | 57 | 4.724 | -6 | -86 | 30 | Occipital_Sup_R | 50 | 5.022 | 28 | -64 | 34 |

| | | | | | | | | | | | |
|---|---|---|---|---|---|---|---|---|---|---|---|
| Frontal_Sup_2_L | 187 | 4.719 | -26 | -8 | 52 | Parietal_Sup_R | 91 | 5.001 | 26 | -68 | 58 |
| Temporal_Inf_L | 279 | 4.711 | -50 | -66 | -6 | Parietal_Sup_R | 91 | 3.897 | 16 | -56 | 72 |
| Rectus_R | 49 | 4.618 | 6 | 44 | -16 | Precuneus_L | 206 | 4.942 | 0 | -70 | 46 |
| Location not in atlas | 137 | 4.521 | -4 | -24 | 28 | Cingulate_Post_R | 101 | 4.890 | 6 | -42 | 22 |
| Cingulate_Post_L | 137 | 3.978 | -4 | -46 | 30 | Temporal_Sup_R | 62 | 4.610 | 46 | -28 | 0 |
| SupraMarginal_L | 74 | 4.407 | -50 | -38 | 24 | Parietal_Inf_L | 39 | 4.584 | -44 | -26 | 36 |
| Occipital_Inf_R | 36 | 4.327 | 32 | -88 | -12 | Supp_Motor_Area_R | 117 | 4.492 | 10 | -12 | 52 |
| Cerebelum_6_R | 36 | 4.127 | 30 | -74 | -22 | | | | | | |

**Figure S1: Comparison of entropy across scales between adults and children**

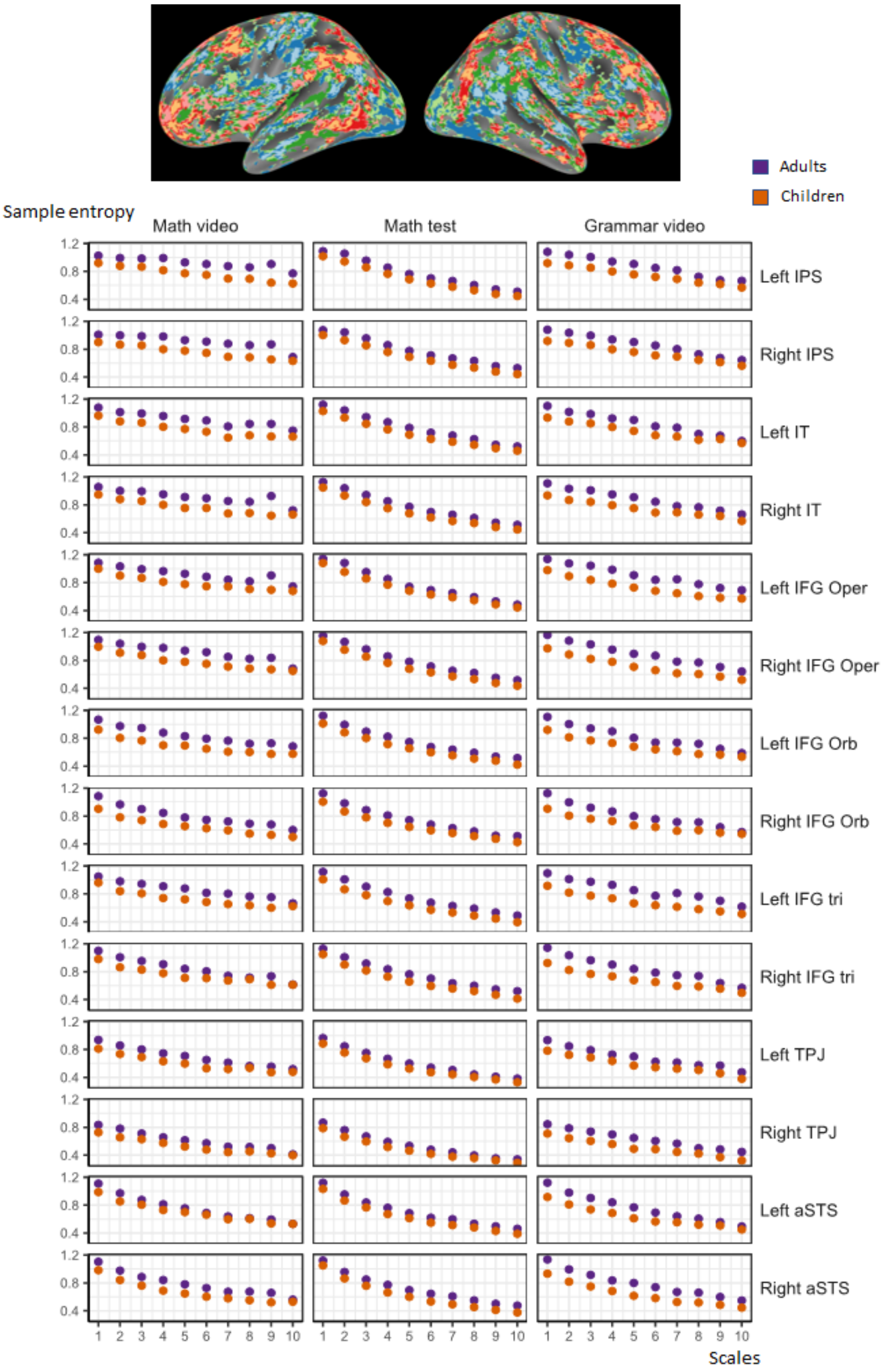


**Top**: Whole brain map showing the difference of average entropy across all timescales between adults and children for our three tasks, with the same color code used in Figure 3. **Bottom**: Mean entropy at each timescale (1 to 10) for the math video (left), the controlled math task (middle), and the grammar video (right), in adults (purple) and children (orange), in 14 regions of interest.

We note that neural entropy at short timescales was higher for adults compared to children in all regions of interest and for all tasks. In particular, after averaging over timescales 1 to 5, all t-tests ps were lower than 0.05, and 12/42 were below 0.0036, i.e. the threshold after Bonferroni correction for multiple comparisons in 14 ROIs. However, this difference faded at longer timescales. After averaging timescales 6 to 10, we noted that the difference between adults and children was mostly significant in math-related regions for the math tasks (all ps < 0.03; math video in the left IPS being the only one for which p was below the threshold after Bonferroni correction), and in language-related regions for the grammar task (all ps < 0.03, except in the left aSTS, p below the threshold after Bonferroni correction in 3 ROIs).

***Figure S2. Correlation between neural maturity and neural entropy at short and long timescales for both video and grammar video lessons***

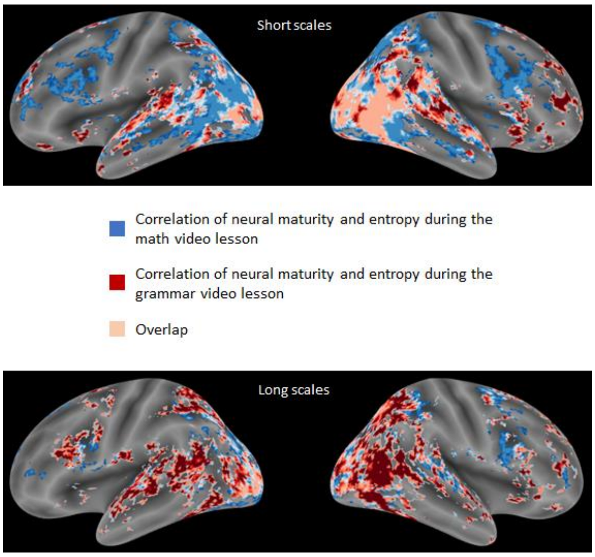


***Figure S3. Correlations and entropy are complementary measures of the signal temporal properties***

Top: two highly correlated periodic signals with different entropy values. Middle: two uncorrelated periodic signals with similarly low entropy values. Top: two uncorrelated signals with similarly high entropy values.

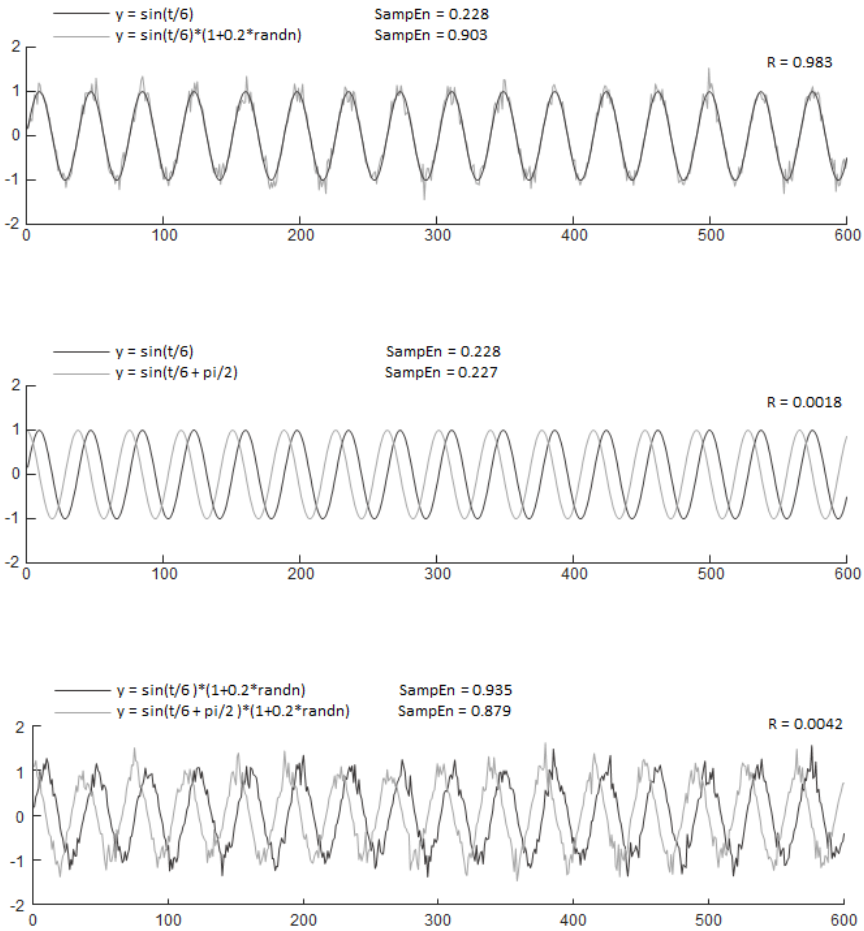
